# KuPID: Kmer-based Upstream Preprocessing of Long Reads for Isoform Discovery

**DOI:** 10.64898/2025.12.05.692400

**Authors:** Molly Borowiak, Yun William Yu

## Abstract

Eukaryotic genes can encode multiple protein isoforms based on alternative splicing of their transcribed regions. Most modern novel isoform discovery methods function by identifying and assembling exon splice junctions from an RNAseq sample. However, splice junctions can only be accurately annotated with time-intensive dynamic programming alignment. This manuscript introduces KuPID, a method for preprocessing long RNAseq reads with the goal of better identifying novel isoform transcripts. KuPID utilizes kmer sketching as a pre-filter to quickly pseudo-align reads to known reference isoforms. Full alignment need only then be applied to reads that are most relevant to isoform discovery. Not only does KuPID speed up the discovery pipeline, it also increases downstream accuracy by filtering out extraneous reads. KuPID preprocessing simultaneously increases the f1 accuracy of isoform discovery pipelines by up to 16.7 points while decreasing the runtime by a factor of 2-3x. An optional mode permits a KuPID sample to be paired with both isoform discovery and transcript quantification. Code availability: https://github.com/mboro2000/KuPID.git

## 1 Introduction

Transcription is the process by which DNA segments are copied and synthesized into proteins. Complex regulatory mechanisms enrich the size and diversity of the human transcriptome. The DNA sequence for a eukaryotic gene is composed of both protein-coding (exon) and non-coding (intron) regions. When transcribing a gene, pre-messenger RNAs are spliced to remove the gene’s introns and ligate its exons into the final transcript. Alternative splicing (AS) enables that same gene to encode multiple alternative protein isoforms. AS events can involve intron retention, exon skipping, or alternative splice sites within the exon [1]. Changes in transcriptional promoter usage and polyadenylation further increases the amount of isoform diversity [2].

As of now, more than 95% of human genes have been found to undergo alternative splicing [3]. Novel isoform discovery methods aim to detect and annotate alternative transcripts to complete the expanding human transcriptome. By doing so, they can also grant insight into the mechanisms of various biological functions. Alternative splicing events have been found to have well-established roles in processes ranging from cell differentiation and survival, to responses to stress [4]. In addition, aberrant splicing has been associated with multiple disease-causing single nucleotide polymorphisms [5]. Isoform discovery (ID) is a common feature of modern transcript quantifiers, which measure changes in isoform abundances through high-throughput RNA sequencing (RNAseq). [6–8] This pairing enables analysis of how the expression of alternative isoforms can differ in response to conditions such as cell state and disease, granting further insight into their function [9].

Despite their importance, it can be difficult to detect many functionally relevant novel isoforms. Unlike constitutively spliced transcripts, alternative transcripts often exhibit unique temporal or cell-type specific expression patterns [10]. Without prior knowledge of their required underlying conditions, such isoforms might rarely appear in an RNAseq sample. Furthermore, many discovery methods employ read support thresholds when evaluating potential novel isoforms [7, 11]. If the rare isoforms are expressed at low levels, ID methods could still fail to detect them in the samples where they are present. Accurate isoform discovery methods are vital to ensure that context-specific isoforms can be studied.

The recent rise of 3rd-gen RNA sequencing has been vital for isoform discovery. The short reads produced by 2nd-gen sequencing are highly accurate, but can omit the ordering of exons and introns in the original transcript [8]. The PacBio and Oxford Nanopore (ONT) platforms can instead sequence full-length transcripts, making it easier to accurately assemble the final isoform [12–14]. Key inefficiencies remain in modern discovery pipelines even with the use of long transcript reads. Current methods require all RNAseq reads to be aligned to a reference genome. The splice junctions of the reads are then identified and assembled to detect novel isoforms [7, 11, 15]. Unfortunately, alignment is time-intensive. The main output of isoform discovery is the exon annotations of the newly found transcripts. For this aim, only the alignment of reads transcribed from novel isoforms should be necessary. However, we typically cannot tell which reads are novel without some form of reference alignment. Given the size of the average RNAseq experiment, isoform discovery methods end up processing millions of query reads that are irrelevant for discovery.

In this manuscript, we introduce Kmer-based Upstream Preprocessing for Isoform Discovery, or KuPID. KuPID reduces runtime by filtering RNAseq reads to those likely to represent novel transcripts so only they need to be mapped. Surprisingly, despite being a lossy filtering step, KuPID actually improves downstream accuracy for the ID pipelines it’s paired with. Instead of trading off one for the other, KuPID provides gains in both speed and accuracy.

## 2 Algorithm Overview

KuPID processes long RNA read sequences exclusively. It requires a reference transcriptome formatted as the sequences of all known transcripts. In its main ‘discovery’ mode, KuPID will output a subset of the reads that are most likely to be sequenced from a novel isoform. The read subset can be submitted to the isoform discovery method of the user’s choice. KuPID consists of three main stages:

1. **Kmer sketching**: We create simplified representations of the RNAseq reads and reference transcriptome that will enable fast comparisons between the two.
2. **Pseudo-alignment to reference transcriptome**: Each RNAseq read is pseudo-aligned with a sparse chaining procedure that finds the largest set of colinear kmer matches between the read and the reference transcriptome.
3. **Read Selection**: We use the sparse chaining results to gauge if any of the reads deviate from the set of known isoforms. If there are substantial gaps in the pseudo-alignment, KuPID concludes that it is likely transcribed from a novel isoform.

KuPID offers an optional mode for end-users that also want to quantify the transcripts in their RNAseq sample. In its ‘quantify’ mode, KuPID will output a subsample of the reads that map to each annotated isoform in addition to the likely novel reads. This mode speeds up alignment, but does not prioritize improvements in isoform discovery.

### 2.1 Kmer Sketching via FracMinHash

Kmer sketching methods are a class of randomized algorithms that focus on constructing smaller and simplified representations of string-formatted data. When applied to large datasets with complex tasks, such as large-scale sequence comparison, sketching offers improved storage and runtime efficiency [17]. We convert every RNAseq read and reference transcript into a subset of its representative kmers using the FracMinHash sketch method [18]. We define *Σ* = {*A, C, G, T*}, and let *Σ*^*k*^ represent the set of nucleotide kmers, or k-length substrings. A hash function *h*: *Σ*^*k*^ → [1, *M*] maps every kmer in the sequence to a numerical value. In our implementation, we apply the invertible integer hash function from minimap [19]. The kmer sketch for a sequence *S* with a set of kmers *W* ⊂ *Σ*^*k*^ is given as 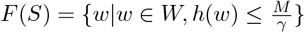, where *γ* determines the fraction of kmers chosen.

### 2.2 Pseudo-alignment to Reference Transcriptome

We first determine a limited reference space *S*(*q*) by the set of isoforms that share at least 1 sketched kmer with the mapping query *q*. We define *R* as the set of all reference isoforms, *Q* as the set of all RNAseq queries, and *Σ*^*k*^ as the set of all potential kmers. To quickly find *S*(*q*), we use a separate data structure *κ*: *Σ*^*k*^ → 𝒫 (*R*) that maps each sketched kmer to the isoforms it’s found in. For a given kmer *x, κ*(*x*) = {*r*|*x* ∈ *F* (*r*)}. A size limit of |*κ*(*x*)| ≤ *b* is placed to remove common kmers and to curtail the size of the search space. We can then formally define the search space *S*(*q*) by 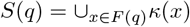.

We narrow the search space further by selecting all isoforms from *S*(*q*) that have the maximum number of unique kmer matches with the query. We define the set of selected isoforms as 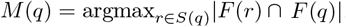. We then create an anchor table *A*_*qr*_ of exact kmer matches between the query *q* and each of the selected isoforms *r* ∈ *M* (*q*). Each anchor *x*_*i*_ = (*x*_1*i*_, *x*_2*i*_) represents a match between the kmer at position *x*_1*i*_ on *q* and position *x*_2*i*_ on *r*. We apply a dynamic programming algorithm to determine the optimal chain, or subsequence of colinear anchors. Every anchor in the optimal chain 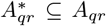 is ordered and strictly increasing so that 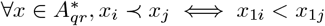 and *x*_2*i*_ *< x*_2*j*_ [20].

We define our optimal chaining score as the total number of anchors in the ordering, and prioritize chains that incorporate as many of the anchors from *A*_*qr*_ as possible. We define 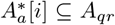 as the optimal chain up to the ith anchor in *A*_*qr*_ such that *x*_*i*_ is included in the chain, and 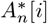as the optimal chain where *x*_*i*_ is not included. We let *f*_*a*_[*i*] and *f*_*n*_[*i*] be the respective optimal chain scores up to the ith anchor. Throughout the chaining process we track *FP*_*l*_[*i*] and *LP*_*l*_[*i*] ∀*l* ∈ {*a, n*}, the first and last anchors included in the optimal chain.

Our chaining procedure is noticeably simpler than other anchor-based methods. Anchor-based chaining methods are often used as an intermediate step in seed-chain-extend alignment. After finding the optimal set of colinear anchors for two sequences, dynamic programming (DP) alignment can be extended to the unmatched regions between chains. Existing chaining methods often penalize large gaps between anchors and / or incorporate the amount of anchor overlap into the chain score [20]. These scores are more appropriate for producing an accurate final alignment. However, we found that pseudo-alignment without extension is sufficient to judge if a query is similar to any reference isoform. We also allow large gaps in the optimal chain instead of penalizing them. We define the gap between two adjacent anchors *x*_*i*_ and *x*_*j*_ in the chain as *g* = | (*x*_1*j*_ − *x*_1*i*_) − (*x*_2*j*_ − *x*_2*i*_) |. We monitor *g*_*l*_[*i*] ∀*l* ∈ {*a, n*}, the largest gap in the optimal chain up to the ith anchor. Tracking the presence of gaps allows us to detect discrepancies in the distances between kmer matches that could indicate alternative splicing.

Similar anchor-based chaining methods employ a banded heuristic to limit the runtime of the DP algorithm [21, 22]. With our simplified chaining formula, many of the anchors will chain to the anchor that directly precedes it. The exceptions to this rule will be if the anchors involve an erroneous kmer, or involve kmers repeated multiple times on the query and reference sequences. KuPID does not allow a kmer to be used more than once in the chain, and aims to skip over the repetitive entries. Anchors that map to the same kmer have repeated *x*_1_ or *x*_2_ positions. As such, we use a band width of *m*^∗^ where we evaluate previous DP entries until we have encountered at least *m* unique kmers on both the query and reference sequence.

We fill out the remaining entries in the table as follows (additional details in A.1):

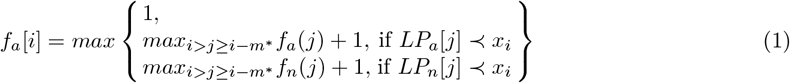

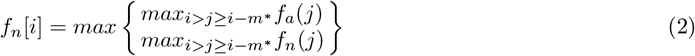

We calculate a similarity score for each RNAseq query based on the Jaccard index, a known metric in gauging overlap between kmer sketches [23]. The Jaccard index corresponds to 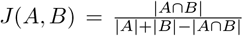, where *A* and *B* are kmer sets from different sequences. We found the Jaccard index was an excellent base metric, but wanted to incorporate the multiplicity and ordering of the kmers in the chain. We define {*F* [*S*]} as the multiset representation of the sketch *F* [*S*]. The multiplicity of a kmer *x* ∈ {*F* [*S*]} reflects the number of times the kmer occurs in the sequence S. The final similarity score for query read *q* is:

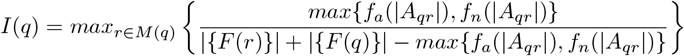

### 2.3 Read Selection

Novel candidate reads are chosen based on the following criteria:

1. Evidence of alternative splicing (AS) events
2. Evidence of mutually exclusive and / or novel exons
3. Evidence of alternative transcription start / stop sites (ATSS)

For a query read *q*, the chaining procedure returns the optimal chaining score and the largest gap *g*^∗^(*q*) in the chain. We define *n* as the expected minimum length of an exon in the reference transcriptome. If *g*^∗^(*q*) > *n*, we infer the read has an AS event and output it as a novel candidate.

We then select the reads that have discernible novel exons. While novel exons can create gaps in the pseudo-alignment, the chaining procedure doesn’t check for reads that either begin or terminate with a novel exon. For each pairwise pseudo-alignment, we calculate the lengths of any unmatched 5’/3’ overhangs along the reference and query sequences.

We define

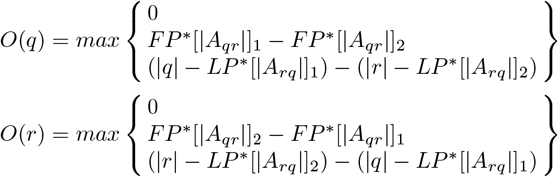

When *O*(*q*) > 0 or *O*(*r*) > 0, there is a discrepancy in the lengths of the exons that lie at the 5’ / 3’ ends of the two sequences. If the discrepancy is sufficiently large, it indicates the presence of a mutually exclusive or novel exon in the query. We automatically select the query read if *O*(*q*) > *n*, as it indicates that the query has at least 1 exon that is not present in the reference transcript. If *O*(*r*) > 0, there are 2 possible scenarios: 1) The query read possesses a shorter novel exon, or 2) the query read was prematurely truncated. We check for truncation by estimating if the query’s unmatched region is sufficiently long enough that it likely contains a novel exon. For 90% of all query reads, the first kmer match will be located within *z* base pairs of the 5’ end, where *z* is solved for with the CDF of the geometric distribution:.9 = 1 − (1 − *s* ∗ (1 − *ke*))^*z*^. The query read will be selected if *FP* ^∗^[|*A*_*rq*_|]_1_ > *z* + *n* or (|*q*| − *LP* ^∗^[ |*A*_*rq*_|]_1_) > *z* + *n*.

KuPID selects additional reads based on the mode selected by the end-user. KuPID offers two modes: ‘discovery’ and ‘quantify’. While the former prioritizes novel isoform discovery alone, the latter can be paired with transcript quantification. In ‘discovery’ mode, KuPID will detect reads with ATSS based on their similarity scores. In theory, the lower the similarity score, the more likely the read is to represent a novel isoform. In practice, it can be difficult to distinguish individual novel and annotated reads from each other. Long sequence reads are frequently truncated due to nonsense mutations, fragmentation during library prep, or RNA degradation [14, 24]. As such, we focus primarily on classifying whole groups of reads that map to a single isoform. The higher the average similarity score of a read group, the more likely it contains only annotated reads. We iteratively add all the reads from the group with the lowest squared average score until we reach the desired number of novel candidates. We limit the size of the candidate set to *c* |*AS*| − |*NE*|, where *AS* is the set of chosen reads with alternative splicing and *NE* is the set of reads with novel exons. *c* is a user-selected parameter that reflects the expected ratio of novel reads without AS to novel reads with AS within the sample.

In ‘quantify’ mode, KuPID will randomly select at most *l* of the reads that map to each known isoform in the RNAseq sample. KuPID will also return a scale factor for each annotated isoform based on the total number of reads that mapped to it. After running the KuPID sample through the quantification method of their choice, a provided script will derive the final quantification results by multiplying the abundance estimates by the scale factors.

## 3 Results

### 3.1 Data Simulation

KuPID was evaluated using a set of simulated PacBio HiFi reads from chr1-22 of the human p14 genome. We created novel isforms via two methods:

1. **YASIM**: An external method that generates isoforms as novel combinations of existing splice junctions [25]
2. **Reduction**: Selecting a random subset of isoforms to treat as novel

We generated 4 novel annotation sets with each method. Based on our chosen complexity index of 9, YASIM automatically generated approximately 43000 isoforms for each annotation set, with a mean of 4.46 isoforms per gene. For the reduction method, we randomly chose 20% of the isoforms in chr1-22 of the human genome to be novel and removed those isoforms from the reference set. The isoforms from the reduction method were filtered so that each gene in the reference retained at least 1 transcript. For both methods, the novel annotation sets were filtered so that every transcript was at least 250 base pairs (bp) long and so there were no intron chain matches with the reference.

We used YASIM to generate the sequencing depth of transcripts within the novel and reference annotation sets. The novel and reference sets each had an average gene depth of 20, and their abundances were generated independently of each other. Multi-pass sequencing of the transcripts was simulated via pbsim3 [26] with a total of 10 passes and an error rate of 10% per run. The multi-pass results were ran through the biopython ccs package to generate HiFi reads with a mean gap-compressed identity of 99.82 [27, 28]. We subsampled the HiFi reads to create datasets with varying percentages of novel reads. Each dataset contained all of the novel isoform reads, and a random subset of the annotated reads.

#### Parameter Selection

For our experiments, we used 22-bp kmers and assumed an error rate of.002. Our sketches used *γ* =.1 of all kmers in the reference transcriptome and RNAseq reads. By default, we set n = 30 as the expected minimum exon length, and b = 16 as the maximum amount of unique isoforms we’d allow a kmer to map to. In addition, we set B =.98 as the expected maximum score of a novel read, and z = 100 as the maximum number of consecutive unmatched bp under the assumption of non-mutually exclusive exons. For ‘quantify’ mode, we set *l* = 5 as the default amount of reads to sample from each annotated isoform. We found that *m* = 3 works well as the band width for our chaining procedure, as it was rare to have erroneous anchor matches. End-users should opt for a larger bandwidth if they have a high error rate or a smaller kmer size. We settled on using a scale factor of *c* = 1.5 when selecting potential novel reads. End-users can increase *c* to improve the method’s sensitivity to isoforms with ATSS. However, overly high values of *c* can hinder the precision of discovery results by allowing additional annotated reads into the sample.

### 3.2 Pipeline Overview

We compared the performance of isoform discovery and quantification methods using the original RNAseq sample or the KuPID-processed reads. All experiments were run on an Intel(R) Xeon(R) Gold 5416S processor with 3 threads. All read sets were first aligned to release 114 of the Ensembl human genome with the minimap2 aligner [22, 29]. The alignment output was submitted to three ID pipelines: IsoQuant, flair, and stringtie2 [7, 11, 15]. Each method employed a reference annotation in guided mode to aid their discovery process. For our reference transcriptome, we used release 48 of the GENCODE human database [30]. The ID pipelines assembled the alignment results into prospective models of the transcripts they believed to be present within the RNAseq sample. The pipelines then outputted an annotation file of the exon features for each transcript model.

### 3.3 KuPID-discovery Enables More Accurate and Efficient Isoform Discovery

We evaluated KuPID’s performance as a novel candidate filter on each of the RNAseq data samples described in Section 3.1. The application of KuPID preprocessing increased the f1 accuracy of all three ID pipelines in comparison to using the original RNAseq sample (Fig 2a). This improvement was consistent across the YASIM and reduction methods of representing the novel transcripts. For the majority of the pipelines, preprocessing with KuPID increased both the precision and recall of the ID results (Fig 2b). We posit that the increase in precision was directly related to KuPID reducing the percentage of annotated reads in the sample (Appendix Table 1). The precision would worsen whenever the discovery pipelines assembled a transcript model that was not actually present in the sample. Any annotated reads could be included in an assembly and lead to the ID method’s hallucinating a false positive. By removing extraneous annotated reads, KuPID curtails the number of false positive models.

**Fig. 1:**
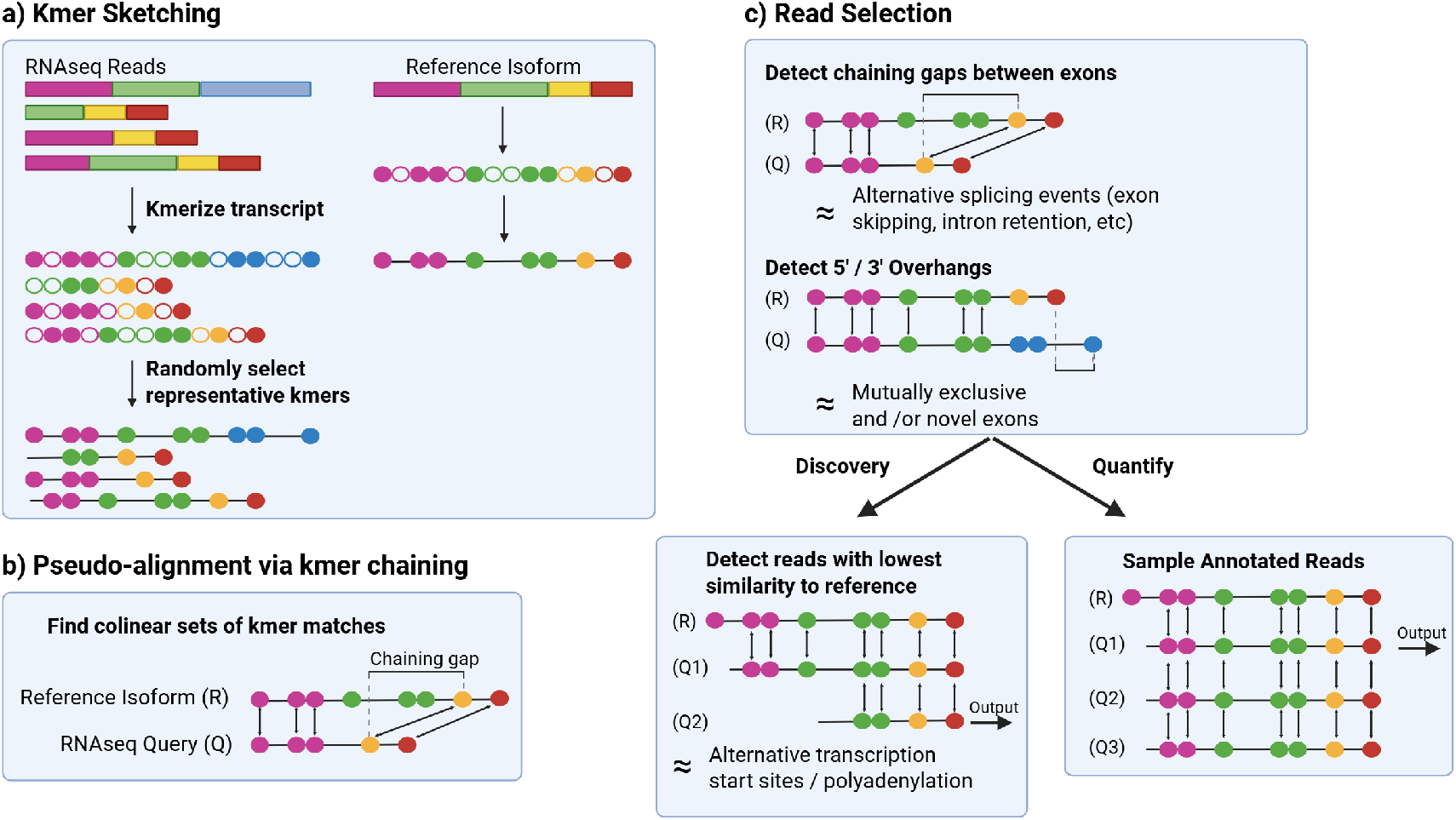
KuPID pipeline for detecting novel isoform reads. Given an RNAseq sample, KuPID returns a fasta of the reads that were likely transcribed from novel isoforms. First, it converts the sample and a reference transcriptome into compact kmer sketches. Then, it finds kmer matches between the sketches to pseudo-align the RNAseq queries to the reference. KuPID determines which reads are novel based on if they have substantial unmatched regions. Given the user’s choice of mode, KuPID can also output a subsample of the reads mapped to each reference isoform for quantification [16]

**Fig. 2:**
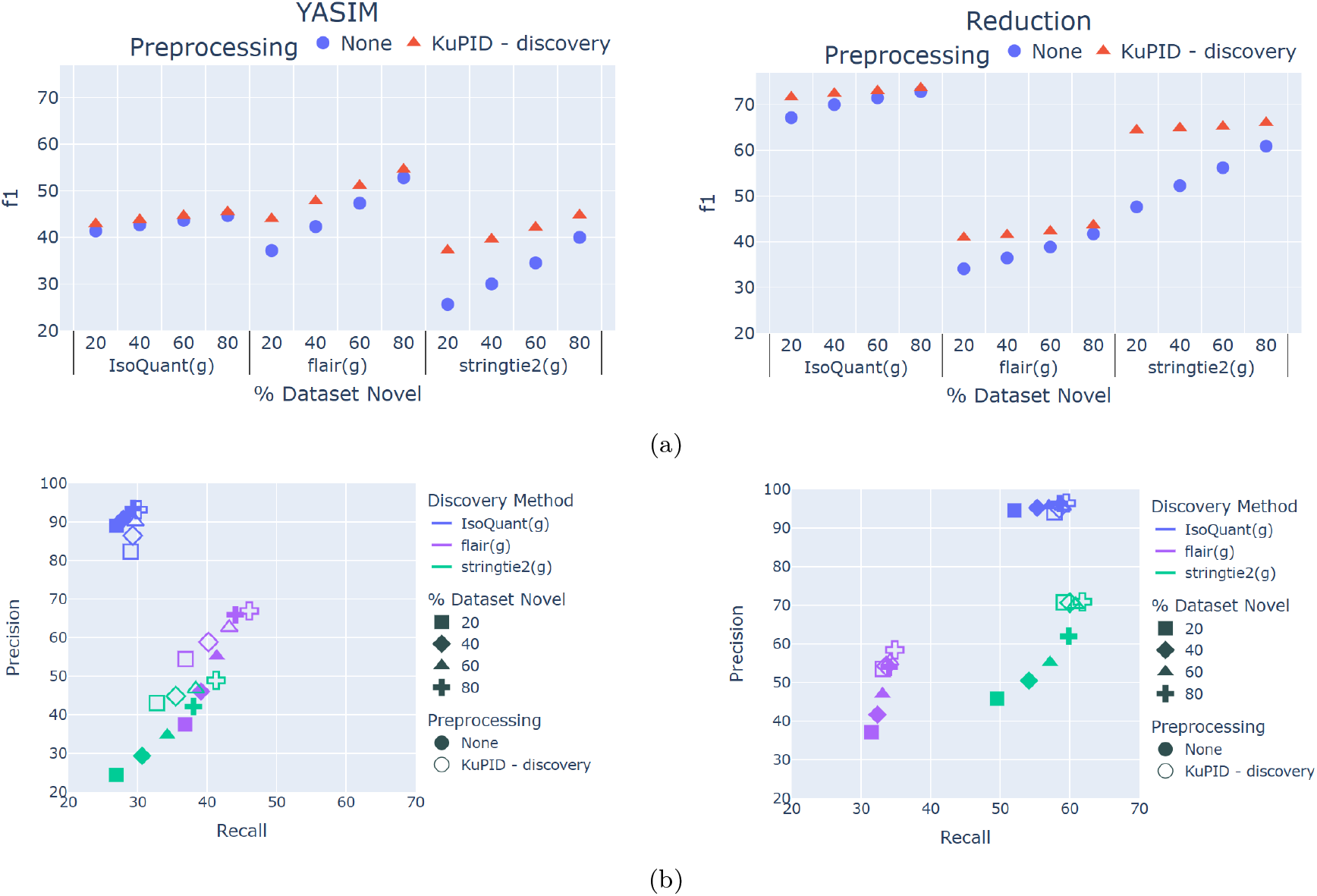
Evaluation of Isoform Discovery Results using KuPID-discovery reads. a) f1 accuracy of the novel transcripts found by the isoform discovery methods. b) Precision and recall of the true novel transcripts found by isoform discovery (ID) methods. The ID methods were applied to either non-processed reads or reads processed with KuPID in ‘discovery’ mode. The percentage of reads from novel transcripts in the original sample ranged from 20-80%. The novel transcripts were generated using either the YASIM (left) or reduction method (right).

KuPID-discovery also minimized alignment bottlenecks by reducing the size of the sample file. For the majority of the datasets, it was faster to run KuPID-discovery and then apply the minimap2 aligner than it was to align the non-processed reads (Fig 3). The exception to this was the extreme case of the dataset containing 80% or more novel reads. The runtime speed-up was most prominent for the datasets that initially contained few novel reads, as they saw the greatest reduction in sample size following the application of KuPID. In the best case scenario where as few as 20% of the initial sample reads were novel, it was at least 2-3x as fast to run the sample through KuPID than to apply full alignment.

**Fig. 3:**
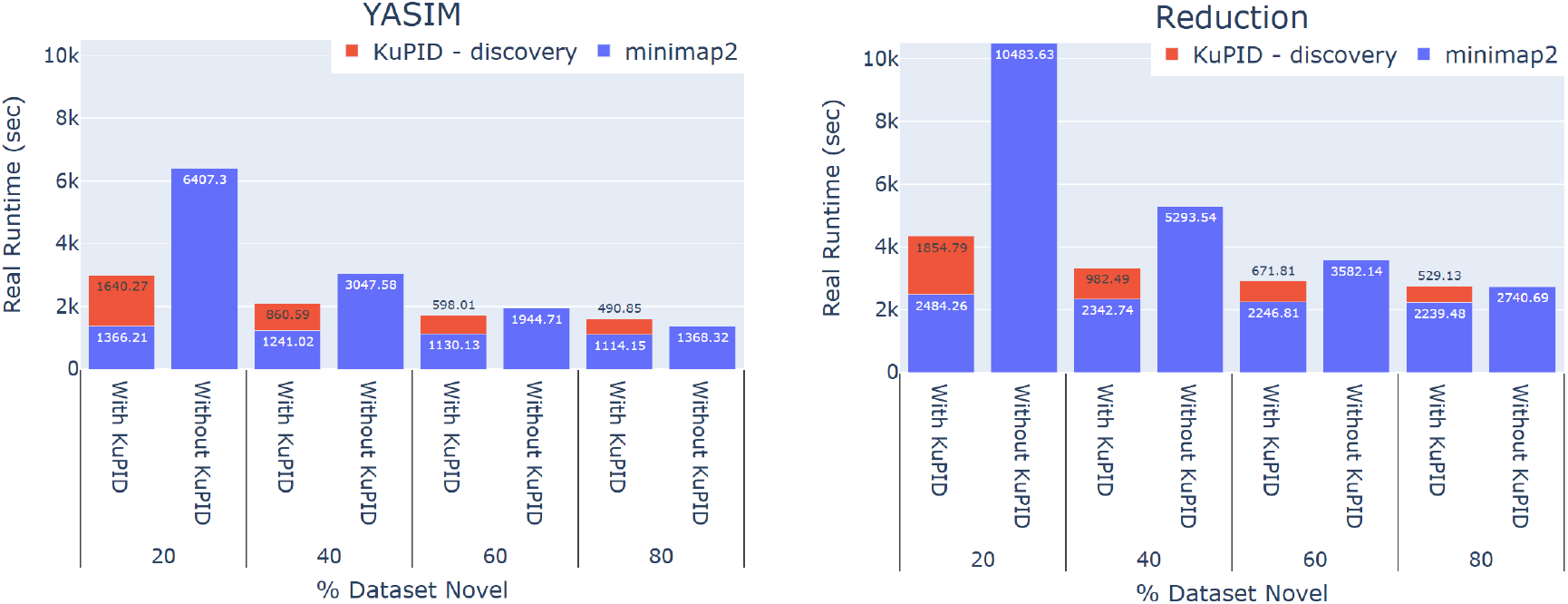
Applying KuPID to RNAseq samples reduces alignment runtime. Average time in seconds needed to align RNAseq reads with the minimap2 software. We compare the runtime of aligning the non-processed reads to the reads selected by KuPID-discovery. The runtime for the KuPID reads encompasses both the time of aligning the reads and the time of applying KuPID to the original sample. The novel transcripts were generated using either the YASIM (left) or reduction method (right).

### 3.4 KuPID-discovery can Detect Difficult Novel Transcripts

Surprisingly, the addition of KuPID to the ID pipelines almost uniformly improved the recall performance (Fig 2b). With KuPID reads, the discovery methods could detect a number of novel transcripts that would otherwise have been hidden from them in the original RNAseq sample. Oddly enough, the blame for the worse recall seems to lie with the annotated reads. When comparing the performance of the non-KuPID samples, the ID methods consistently recalled fewer novel transcripts from samples with higher proportions of annotated reads (Fig 2b). These datasets contained identical novel reads, and only differed in their number of annotated transcripts. Thus, the inclusion of annotated reads in the sample appears to mask novel transcripts from the ID pipelines.

The impact of annotated reads was felt not just across the sample, but particularly at the gene-level. We found that the ID method’s ability to detect novel transcripts was also impacted by the composition of isoforms expressed by a given gene. In our simulated data, any gene that expressed a novel transcript could simultaneously express annotated transcripts as well. Based on the original RNAseq samples, we compared the ability of the ID methods to detect novel transcripts that were expressed alongside annotated reads versus novel transcripts that were not. ID pipelines often had less success in detecting novel isoforms transcribed from genes with mixed expression (Fig 4a). The addition of KuPID to the pipelines mitigated the effects of masking and enabled the discovery of these difficult transcripts. Preprocessing the RNAseq samples increased recall of alternative transcripts at both the gene-(Fig 4b) and sample-wide levels (Fig 2b). The magnitude of KuPID’s impact on recall was not uniform. For IsoQuant and flair, the improvement of recall at the gene-level was somewhat marginal. Of all the pipelines, stringtie2 had the largest increase in the number of novel transcripts detected when paired with KuPID.

**Fig. 4:**
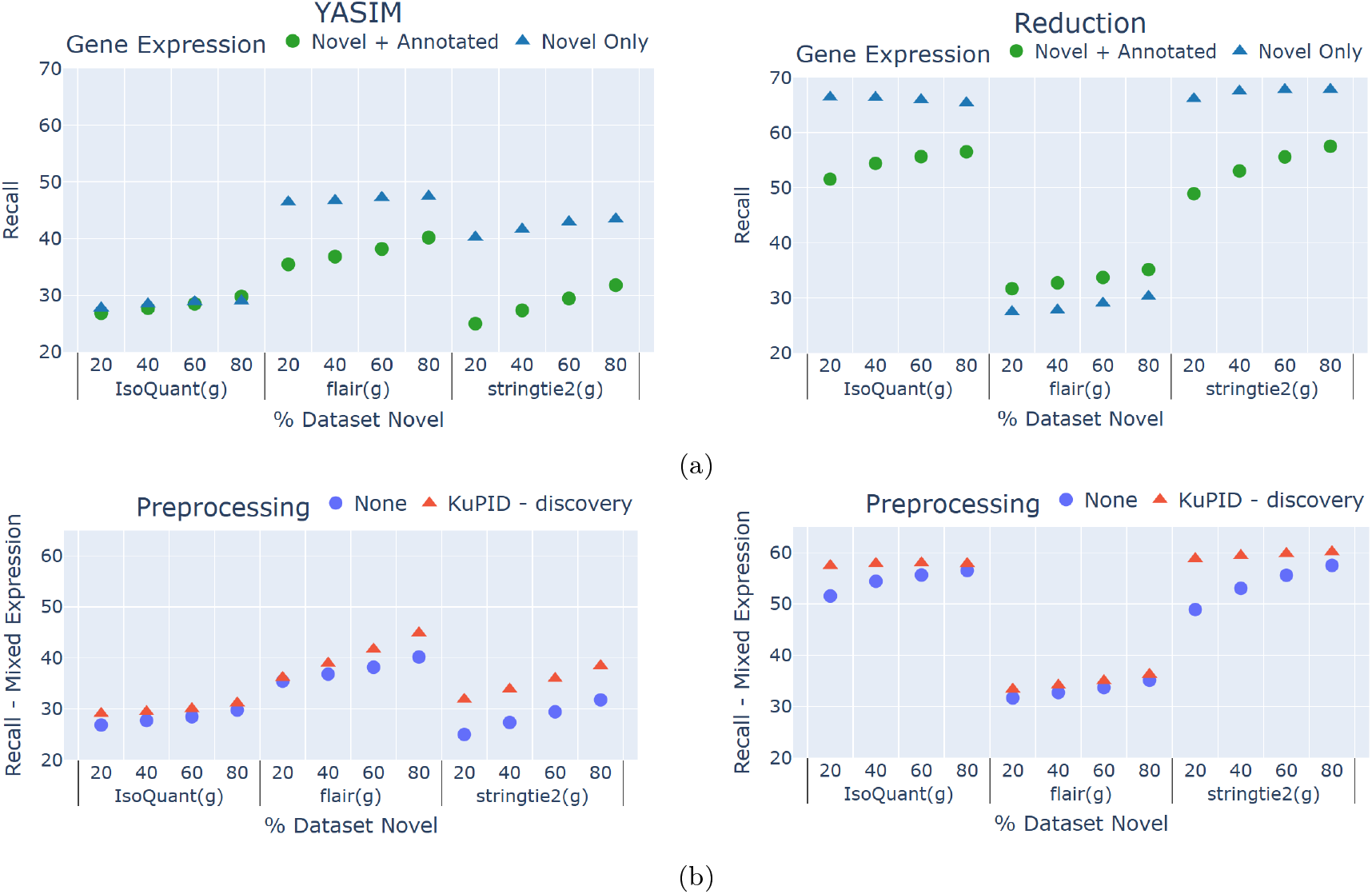
Isoform discovery methods have difficulty in de tecting is oforms fr om ge nes wi th mixed expression. KuPID can detect novel transcripts that are otherwise masked by annotated reads. a) Recall of the true novel transcripts found by isoform discovery (ID) methods when applied to nonprocessed RNAseq samples. Recall is compared between genes that only expressed novel isoforms and genes that expressed both novel and annotated isoforms. b) Recall of the novel isoforms transcribed from genes with mixed expression. The isoform discovery methods were applied to either the non-processed RNAseq sample or the reads selected by KuPID-discovery. The novel transcripts were generated using either the YASIM (left) or reduction method (right).

### 3.5 KuPID-quantify Enables More Efficient Transcript Quantification

KuPID’s main mode (KuPID-discovery) prioritizes the accuracy of isoform discovery methods. KuPID’s optional mode (KuPID-quantify) can accommodate both discovery and quantifcation analysis. To illustrate this, we employed KuPID-quantify to create a candidate set for each of the data samples in Section 3.1. We then applied the discovery and quantification modules of IsoQuant, flair, and stringtie2. We found that the discovery results from the KuPID reads closely mimicked the results from the non-processed reads (Fig 5a). The KuPID reads also performed reasonably well in quantifying the abundance of annotated transcripts. We calculated the Spearman’s rank correlation between the true abundance of each reference isoform (reported in Transcripts per Million) and the predicted abundance. Across the data samples, the Spearman correlation for the KuPID processed reads was at least as strong as the correlation with the non-processed reads (Fig 5b). In terms of runtime, KuPID-quantify had similiar efficiency to KuPID-discovery. KuPID-quantify consistently reduced alignment time for all datasets with fewer than 80% novel reads (Fig 5c).

**Fig. 5:**
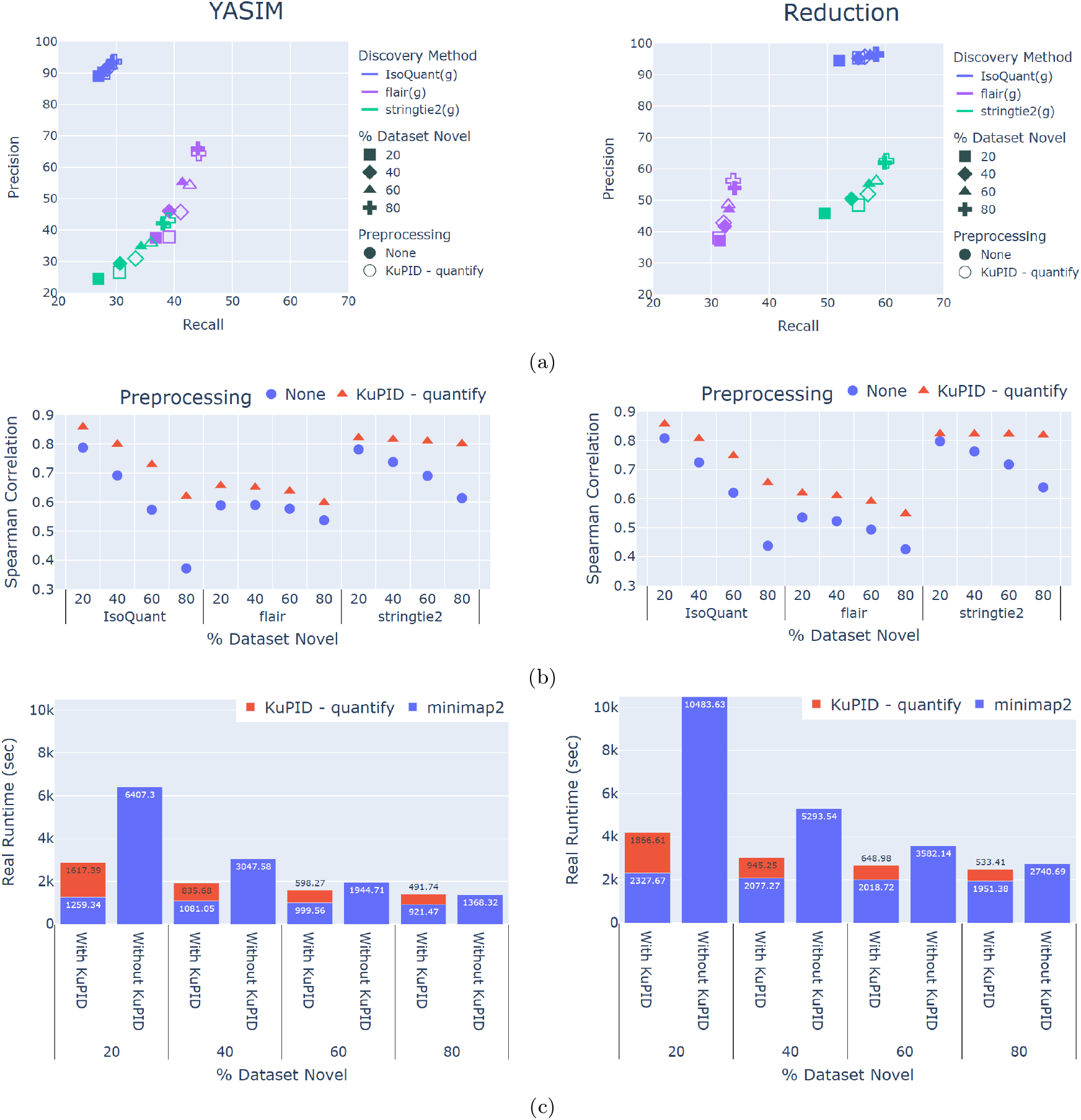
Evaluation of Transcript Discovery and Quantification Performance with KuPID-quantify reads. a) Precision and recall of the true novel transcripts found by isoform discovery (ID) methods. The discovery pipelines were applied to either non-processed RNAseq samples or to the reads selected by KuPID-quantify b) Spearman coefficient of the relationship between the true and predicted abundances of annotated isoforms in the sample. Abundances were reported in Transcripts Per Million. c) Average time in seconds needed to align RNAseq reads with the minimap2 software. We compare the runtime of aligning the non-processed reads to the reads selected by KuPID-quantify. The novel transcripts were generated using either the YASIM (left) or reduction method (right).

## 4 Discussion

Our method KuPID can successfully eliminate long annotated reads from RNAseq samples, directly leading to more efficient and accurate isoform discovery pipelines. KuPID-discovery is most effective when it is applied to data samples where the minority of reads are novel. Even so, KuPID can improve the accuracy of discovery methods irrespective of the total amount of novel reads in the sample.

KuPID is robust to not only the amount of novel reads present, but to the types of novel isoforms as well. We generated novel isoforms with two methods: either creating new isoforms with the external software YASIM, or by randomly reducing the reference transcriptome and treating the removed isoforms as novel. KuPID-discovery successfully improved the accuracy of the ID results of the datasets created with both methods. Notably, the IsoQuant and stringtie2 methods had greater success in recalling novel transcripts from the reduced method than from the YASIM method (Figure 2b). This trend was consistent across both the original RNAseq samples and the KuPID processed reads. One possible explanation could be that the reduction method produced a high number of transcripts with novel exons. Each set of novel isoforms created through reduction had ≈20,000 novel exons, while the YASIM set did not contain any novel exons. This seems to indicate that the presence of novel exons has a strong influence on whether or not a novel isoform will be detected by the pipeline.

Unexpectedly, ID pipelines that pre-processed with KuPID were able to detect a greater amount of the novel transcripts that were present in a sample. This phenomenon likely stems from KuPID reducing read support bias from the prospective transcript models. Besides employing support thresholds, current isoform discovery methods tend to favor transcripts with the greatest amount of read support. The base stringtie algorithm employs a network flow algorithm to reconstruct transcripts [31]. A splice graph is constructed from the sample reads, with each splice path representing a transcript. The algorithm iteratively finds the transcript with the heaviest read coverage before assigning reads to it. IsoQuant uses a similar intron graph algorithm, and removes any transcript with less than 2% read support from the maximum graph coverage [7]. Selecting models with the greatest amount of support is intuitive, but read support can be inflated if a model is covered by both annotated and novel reads. To illustrate this, we found that isoform pipelines have more difficulty in detecting novel transcripts when they are expressed by genes that also express annotated transcripts (Fig 4b). By filtering out annotated reads at the gene-level, KuPID can likely remove read support bias and improve the recall of novel isoforms.

Ensuring that isoform discovery pipelines have sufficiently high recall is vital for detecting context-specific novel transcripts. Alternative transcripts can have unique expression patterns based on a given environment or cell-type [10]. Practically, rare or expensive conditions will have a limited amount of replicates to analyze for novel transcripts. ID pipelines that involve KuPID are able to remove extraneous reads that would otherwise have occluded context-specific transcripts from discovery.

## 5 Conclusion

Our method KuPID can preprocess long RNAseq reads to create an optimal subsample for either novel isoform discovery or transcript quantification. KuPID-quantify can closely approximate discovery and quantification results while decreasing the amount of time needed for downstream analysis. By minimizing read support bias at the gene-level, KuPID-discovery can increase the detection of alternative transcripts that had previously been masked by annotated reads. KuPID-discovery’s success is robust to both the percentage of novel reads in a given sample, and to the variety of alternative splicing present. KuPID can aid in the detection of context-specific transcripts that can only be replicated in relatively few samples.

## Acknowledgments

This research was funded in part by the US National Science Foundation BIO/DBI grant 2531433. We’d like to thank Aidan Zhang from the Yu Lab for his help in developing this manscript.

## Disclosure

The authors have no competing interests to declare that are relevant to the content of this article.

## A Appendix

**Table 1:** KuPID reduces the percentage of annotated reads in an RNAseq sample by an order of at least 3x. Mean percentage of annotated reads present the RNAseq samples. The mean percentage is compared non-processed samples and the samples processed with KuPID-discovery.

**Percent Annotated Reads in RNAseq Sample**
| Novel Isoform Method | No Preprocessing | KuPID-Discovery |
| --- | --- | --- |
| YASIM | 20 | 8.3 |
|  | 40 | 11.51 |
|  | 60 | 16.12 |
|  | 80 | 24.76 |
| Reduction | 20 | 8.65 |
|  | 40 | 11.69 |
|  | 60 | 15.16 |
|  | 80 | 20.57 |

### A.1 Methods

#### Determining Optimal Chains

For each table entry *f*_*n*_[*i*], we define the previous entry that contributed to the optimal chain score as 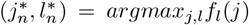. For each table entry, *f*_*a*_[*i*], we define 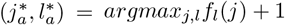 for *x*_*j*_ ≺ *x*_*i*_. We initialize and update the properties of the optimal chain as follows:

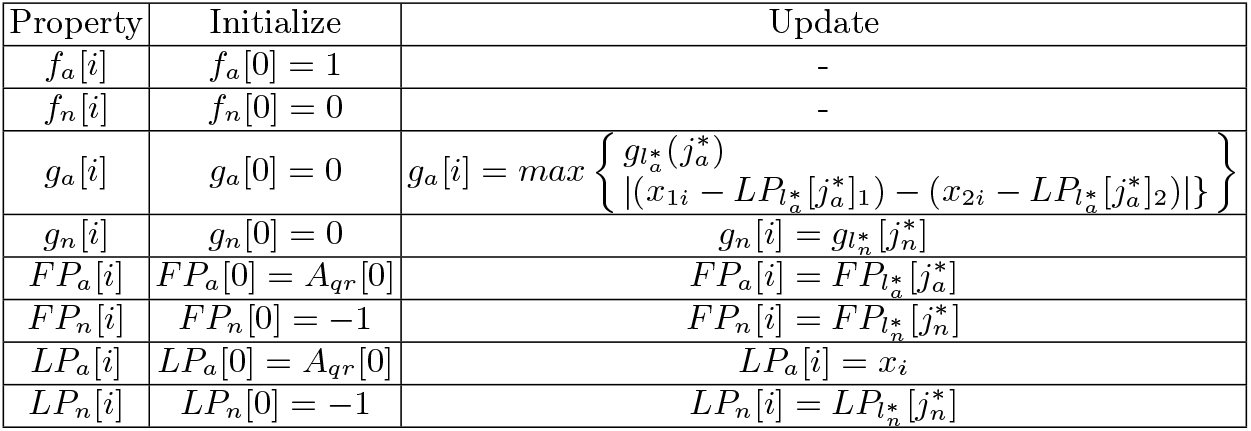

### A.2 Metrics

The ID pipelines were evaluated based on their recall and precision of the novel isoforms generated by YASIM. All accuracy metrics for the annotation files were calculated using gffcompare [32]. The initial annotation files created by the discovery pipelines were processed so that they only included transcripts that were predicted to be novel. To do so, we used gffcompare to compare the ID output to the original set of human chr1-22 annotations. We then omitted every ID annotation that had an intron chain match with the reference set. The recall and precision results were reported from comparing the ‘predicted novel’ annotations to the set of novel transcripts that were present among the HiFi reads. All metrics were reported from the transcript-level and encompassed both exact intron-chain matches and single exon matches.

For our quantification experiments, abundance estimates were reported in Transcripts per Million (TPM). The abundances were calculated relative to the transcript reads that only mapped to known reference isoforms.

## Notes

### Competing Interest Statement

The authors have declared no competing interest.

### Summary of Updates

Minor revisions for grammatical errors and to provide clarity as to why some parameters were used.

